# Human thalamic recordings reveal that epileptic spikes block sleep spindle production during non-rapid eye movement sleep

**DOI:** 10.1101/2023.04.17.537191

**Authors:** Anirudh Wodeyar, Dhinakaran Chinappen, Dimitris Mylonas, Bryan Baxter, Dara S. Manoach, Uri T. Eden, Mark A. Kramer, Catherine J. Chu

## Abstract

In severe epileptic encephalopathies, epileptic activity contributes to progressive cognitive dysfunction. Several epileptic encephalopathies share the trait of spike-wave activation during non-rapid eye movement sleep (EE-SWAS), a state dominated by sleep oscillations known to coordinate offline memory consolidation. How epileptic activity impacts these thalamocortical sleep oscillations has not been directly observed in humans. Using a unique dataset of simultaneous human thalamic and cortical recordings in subjects with and without EE-SWAS, we reconcile prior conflicting observations about how epileptic spikes coordinate with sleep oscillations and provide direct evidence for epileptic spike interference of sleep spindle production. We find that slow oscillations facilitate both epileptic spikes and sleep spindles during stage 2 sleep (N2) at different phases of the slow oscillation. We show that sleep activated cortical epileptic spikes propagate to the thalamus (thalamic spike rate is increased after a cortical spike, p∼0). Thalamic spikes increase the spindle refractory period (p<1.5e-21). In patients with EE-SWAS, the abundance of thalamic spikes result in downregulation of spindles for 30 seconds after each thalamic spike (p=3.4e-11) and decreased overall spindle rate across N2 (p=2e-7). These direct human thalamocortical observations identify a novel mechanism through which epileptiform spikes could impact cognitive function, wherein sleep-activated epileptic spikes inhibit thalamic sleep spindles in epileptic encephalopathy.

## Introduction

Cognitive dysfunction is a common comorbidity in epilepsy (Taylor et al., 2010; Taylor & Baker, 2010; Wickens et al., 2017), but the mechanism remains unknown (Beenhakker & Huguenard, 2009). In the most severe form, cognitive dysfunction is described as an epileptic encephalopathy, where patients have a reduced rate-of-development, plateau, or frank regression in cognitive functions concurrent with the development of epileptiform activity (Specchio et al., 2022). Several epileptic encephalopathies share the trait of interictal epileptic spikes that are potentiated during non-REM sleep, termed epileptic encephalopathy associated with spike-wave activation during sleep (EE-SWAS). In these cases, epileptic spikes during sleep are thought to be mechanistically related to cognitive decline, however existing studies have found that epileptic spike rates in sleep fail to predict cognitive symptoms in EE-SWAS (Binnie, 2003; Bjørnæs et al., 2013; Henin et al., 2021; Larsson et al., 2012).

Non-REM (non-rapid eye movement) stage N2 sleep is characterized by sleep spindles, a 9-15 Hz oscillation implicated in sleep-dependent memory consolidation (Denis et al., 2021; Gais et al., 2002; Latchoumane et al., 2017) and correlated with cognitive measures (M. Hahn et al., 2019; Reynolds et al., 2018). Sleep spindles originate in the thalamus and propagate to the cortex and occur with an approximate period of 2-5 s, dictated by an after-depolarization spindle refractory period (Bal & McCormick, 1996; Fernandez & Lüthi, 2020). Spindles are frequently nested in the up-state of slow oscillations – 0.5-2 Hz oscillations – (Mak-McCully et al., 2017; Schreiner et al., 2022; Steriade et al., 1993) and, critically, may coordinate hippocampal ripples to support memory consolidation (Latchoumane et al., 2017; Staresina et al., 2015).

Epileptic spikes, in contrast to sleep spindles, are cortically-driven pathological events, reflecting summated, excessively synchronous neural activity (Steriade, 2006; Traub & Wong, 1982). Observations in animals (Steriade & Contreras, 1998) and in humans (Velasco et al., 1991, 2002) demonstrate that cortical epileptic spike activity may propagate to the thalamus (Gadot et al., 2022). Epileptic spikes have been proposed to hijack the circuitry that produces spindles and thereby disrupt sleep-dependent memory consolidation (Beenhakker & Huguenard, 2009; Kramer et al., 2021). However, competing observations suggest that epileptic spikes precede – and therefore may induce – sleep spindles, with potentially pathologic consequences for cognitive function (Dahal et al., 2019; Gelinas et al., 2016; Sákovics et al., 2022). How and whether cortical spikes interact and interfere with thalamic sleep spindles in humans is unknown. Direct observations of these dynamics are challenging due to the scarcity of recordings available from the human thalamus.

Using a unique dataset of simultaneous human thalamic and cortical recordings in subjects with and without EE-SWAS, we reconcile prior conflicting observations and provide direct evidence for epileptic spike interference of sleep spindle production. We find that cortical slow oscillations facilitate both epileptic spikes and spindles during N2 sleep at different phases of the slow oscillation. Separately, sleep activated cortical epileptic spikes that propagate to the thalamus inhibit spindle production, most prominently in subjects with EE-SWAS. Given this evidence, we propose the disruption of thalamic spindles by cortical spikes during non-REM sleep as a mechanism of cognitive dysfunction in epilepsy.

## Methods

### Subject data collection

Subjects with simultaneous thalamic and cortical local field potentials and scalp EEG recording collected as part of their clinical evaluation for drug resistant epilepsy between 07/2020 and 11/2021 at Massachusetts General Hospital were evaluated. To ensure appropriate thalamic lead placement relative to the cortical irritative zone, only those subjects found to have instantaneous ictal propagation to the thalamic target were included, resulting in datasets from 9 subjects – thalamic leads targeted the centromedian nucleus in 7 subjects and the anterior nucleus in 2 subjects (see Table 1). Clinical diagnosis, the location of the seizure onset zone, antiseizure medications, and demographic information were collected from chart review (see Table 1). Pre- and post-operative high-resolution MRIs were collected for electrode co-registration. This study was approved by the Massachusetts General Hospital Institutional Review Board.

### EEG data collection, pre-processing, and channel selection

Intracranial and scalp EEG data were collected using the clinical Natus Quantum system (Natus Neurology Inc., Middleton, WI, USA). Depth electrodes (PMT Depthalon depth platinum electrodes with 3.5 mm spacing, 2 mm contacts, and 0.8 mm diameter; or Ad-tech depth platinum electrodes with 5-8 mm spacing, 2.41 mm contact size, and 1.12 mm diameter) were placed in the regions of clinical interest and sampled at 1024 Hz (8 subjects) or 2048 Hz (1 subject, subsequently downsampled to 1024 Hz). Twenty-four hours of scalp EEG data (21 scalp electrodes plus electrocardiogram and electroculogram electrodes) were sleep-staged by a board-certified clinical neurophysiologist (CJC) following standard visual criteria (Grigg et al., 2007; Iber et al., 2007). To control for variability across sleep stages and focus on sleep-activated epileptic spike activity and sleep spindles, we selected only non-rapid eye movement stage 2 sleep segments (N2) for analysis. All N2 sleep segments over the course of the 24-hour interval were concatenated for analysis.

For epileptic spike and spindle detections we evaluated three voltage time series from each subject: 1) Adjacent thalamic depth electrode contacts with the highest amplitude signal in a bipolar reference; 2) Adjacent cortical depth electrode contacts in the clinically determined seizure onset zone in a bipolar reference; 3) Scalp EEG in a bipolar reference CZ-PZ. To detect slow oscillations, we analyzed the scalp CZ contact using a far-field non-cephalic reference placed over the second cortical spinous process (Schreiner et al., 2022; Staresina et al., 2015). The channel and reference used to test each hypothesis are stated in the statistical analysis section below.

### Automatic event detection

#### Epileptic spike detection

To detect epileptic spikes, we extended the method in (Dahal et al., 2019; Gelinas et al., 2016). First, we bandpass filtered the data both forwards and backwards using a finite impulse response (FIR) filter between 25 to 80 Hz (MATLAB function firfilt). We then applied a Hilbert transform, calculated the analytic signal, and the amplitude envelope of this signal. For each voltage signal, the moments when the amplitude envelope exceeded three times the mean amplitude were identified as candidate spikes. To ensure the candidate spikes were not due to gamma-band oscillatory bursts, we also calculated the regularity of oscillations in an interval (±0.25 s) around each candidate spike. To assess the regularity of the signal in this interval, we computed the Fano factor (Eden & Kramer, 2010) estimated by: (i) detrending the interval of unfiltered data at each candidate spike, (ii) identifying peaks and troughs, (iii) calculating the inter-peak and inter-trough intervals, and (iv) estimating the ratio of the variance of the inter-peak and inter-trough intervals to the mean of the inter-peak and inter-trough intervals. We then removed candidates spikes if the maximum amplitude (calculated by rectifying the data and identifying the maximum voltage) of the unfiltered data at the candidate spike was below three times the mean amplitude or if the Fano factor was less than 2.5; we note that Fano factors below 1 indicate a more regular rhythm, with 0 indicating no variability in the inter-peak or inter-trough intervals. Of the resulting spikes, those detected within 20 ms of one another were merged into a single spike detection. We visually confirmed by examining individual detections and the averaged spike waveform that the method accurately detected epileptic spikes in both cortical and thalamic recordings.

#### Spindle detection

We applied an existing spindle detector with robustness to epileptic spikes and sharp transients (Kramer et al., 2021). Briefly, the spindle detector estimates a latent state – the probability of a spindle – using three features: the Fano factor (estimated for data FIR filtered between 3 – 25 Hz), normalized power in the spindle band (9 – 15 Hz), and normalized power in the theta band (4 – 8 Hz). Distributions of expected values for these parameters were determined using manually detected spindles in the scalp EEG of subjects with sleep-activated spikes. To avoid misidentifying spikes as spindles, we applied a cubic spline to the ±50 ms interval around each detected spike before applying the spindle detector (as recommended in Klinzing et al., 2021). Doing so further improves the ability of the spindle detector to reject spikes and accurately identify the beginning and end of spindles. The spindle detector estimates the probability of a spindle in 0.5 s intervals (0.4 s overlap). We detected a spindle when the probability crossed 0.95, chosen by optimizing the sensitivity and specificity of the detector (Kramer et al., 2021). We retained spindles with a minimum duration of 0.5 s, a maximum duration of 5 s, and separated by at least 0.5 s; spindles within 0.5 s of one another were merged into a single spindle detection. We visually confirmed that the method accurately detected sleep spindles in both scalp EEG and thalamic recordings.

#### Slow oscillation detection

We applied the algorithm described in (Mölle & Born, 2011) to detect slow oscillations. Briefly, slow oscillations were detected by: (i) filtering the data into the 0.4 Hz to 4 Hz band using an FIR filter (4092 order); (ii) identifying all negative to positive zero-crossings and the time interval *t* to the subsequent positive to negative zero-crossings; (iii) retaining all intervals of duration 0.5 s ≤ *t* ≤ 2 s (corresponding to oscillations 0.5 to 2 Hz) and identifying the negative peak and peak-to-peak amplitude; (iv) omitting intervals in the bottom 75^th^ percentile of peak-to-peak amplitude and the bottom 25^th^ percentile of the negative peak amplitude; (v) retaining any remaining intervals as slow oscillations.

### Statistical analysis

#### Point process models for slow oscillations, spikes, and spindles

To test hypotheses about relationships between slow oscillations, epileptic spikes, and thalamic or scalp spindles, we first represented the event detections as a sequence of discrete events or point process time series. These stochastic point process time series are completely characterized by the conditional intensity function, representing the history-dependent generalization of the rate function of a Poisson process,

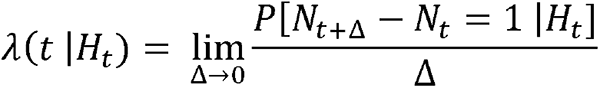

where *P* is a conditional probability, *N_t_* denotes the number of events counted in the time interval (*0,t*] and *H_t_* includes the event history up to time *t*, and other covariates (Truccolo et al., 2005). The logarithm of the conditional intensity, when considering a time discretized point process, can be expressed as a linear function of covariates,

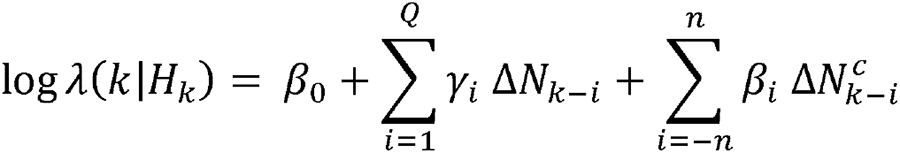

where *k* is the *k^th^* interval of discretized time (with *k* > *Q* and *k* > n), the first term (β_0_) epresents a baseline event rate, the second term represents a self-history autoregressive process, and the third term represents past and future contributions from other covariates. The expressions 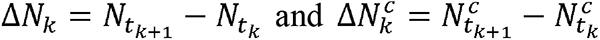 indicate binary time series of increments in event counts, and (β_i_,γ_i_) are parameters to estimate. This discrete-time point process likelihood function is equivalent to the likelihood of a generalized linear model (GLM) under a Poisson distribution and log link function (Truccolo et al., 2005). We estimated model parameters using the fitglm function in MATLAB.

#### Modeling slow oscillations, cortical spikes, and thalamic or scalp spindles

To characterize the relationship between scalp slow oscillations and cortical epileptic spikes we divided time into 0.125 s intervals and created two binary time series. In the first time series, a value of 1 was assigned to intervals containing the downstate of a slow oscillation, and 0 otherwise. In the second time series, a value of 1 was assigned to intervals containing the maximum amplitude of an epileptic spike, and 0 otherwise. When more than one epileptic spike occurred in the same 0.125 s interval, the value of that interval was set to the number of spikes that occurred. Using these point process time series, we estimated the model:

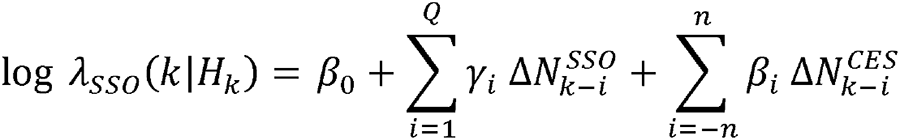

where λ_sso_(*k*|*H_k_*) represents the conditional intensity or conditional event rate of scalp slow oscillations (SSO) given the history of SSO events and the occurrence of cortical epileptic spikes (CES) at past and future times.

In the same way, to characterize the relationship between scalp slow oscillations and thalamic spindle oscillations, we created point process times series and estimated the conditional event rate of scalp slow oscillations. For spindle detections, we assigned each 0.125 s interval a value of 1 if the maximum amplitude of a spindle occurred, or 0 otherwise. Slow oscillations and spindle oscillations were modeled using:

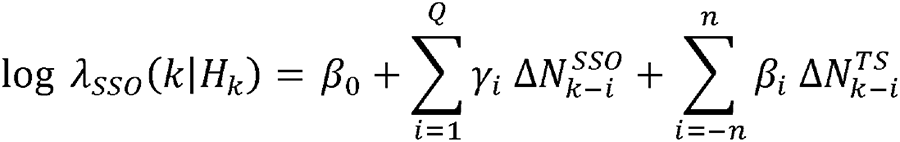

where λ_sso_(*k*|*H_k_*) represents the conditional intensity or conditional event rate of scalp slow oscillations (SSO) given the history of SSO events and the occurrence of thalamic spindles (TS) at past and future times. We applied the same model to assess the relationship between slow oscillations and scalp spindles by replacing 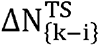 with 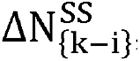, the history and future of scalp spindle occurrences.

Using these models, we estimated the phase relationships between slow oscillations, spikes, and spindles. We note that the prediction of the scalp slow oscillation downstate using cortical spikes and thalamic spindles is not causal. We chose this model structure to assess the temporal relationship of spikes and spindles to the slow oscillation downstate in a manner concordant with past work (Frauscher et al., 2015).

For all models, we tested the model improvement relative to a nested model with only the history of the scalp slow oscillations. To estimate parameter significance within a model, we applied the Wald test. Confidence intervals for the parameters were estimated from the model fits using the function fitglm in MATLAB. Finally, we tested hypotheses across groups of parameters (see Chapter 5 of Greene, 2017) using an F-test implemented in the coefTest function in MATLAB.

#### Procedures to estimate cortical spike propagation to thalamus

To test the hypothesis that cortical epileptic spikes propagate to the thalamus, we estimated: (1) the average thalamic evoked response, (2) an unnormalized cross-correlation histogram, and (3) the conditional event rate of thalamic spikes given cortical spikes. We define each measure here.

The average evoked response assumes a temporally locked relationship between cortical epileptic spikes and thalamic epileptic spikes. To assess this relationship, for each subject, we extracted from the thalamic voltage signal ±1 s intervals centered on the times of each cortical spike. We then estimated the mean thalamic response and standard error across all subjects and cortical spikes and tested the results against overlap with zero. Confidence limits were Bonferroni corrected for multiple comparisons.

To estimate the unnormalized cross-correlation histogram (Harrison et al., 2013), we first created binary time series using 1 to demarcate time-points with the maximum amplitude of an epileptic spike and 0 otherwise. Then we: (i) identified the time *t_i_* each event *i* from the cortical binary time series; (ii) identified the delays of all events in the thalamic binary time series in the interval ±1 s around each *t_i_*; (iii) constructed a histogram from the temporal delays of thalamic spikes relative to the cortical spikes using ≈35 ms bins. To assess the significance of the cross-correlation histograms, we created a null distribution of values by shuffling the inter-event intervals (i.e., time between events) of the cortical binary time series and then re-estimating the cross-correlation histograms. This process accounts for baseline rates of cortical spike occurrence and the distribution of inter-event intervals. We repeated this procedure 5000 times for each subject. Using the shuffled cross-correlation histograms, we created 95% confidence intervals for the null distribution and corrected for multiple comparisons (due to the 60 bins, each of ≈35 ms width) using Bonferroni correction. From the cross-correlation histogram, we computed the number of thalamic spikes in each time bin between [0,1] s after a cortical spike for each patient. To compare the mean number of thalamic spikes in the [0,1] s interval between groups (EE-SWAS, SWAS, and non-SWAS), we modeled the mean number of thalamic spikes with an indicator for group (generalized linear model with log-link and a Gamma distributed response). We compared the mean number of spikes between the EE-SWAS and non-SWAS groups, and between SWAS and non-SWAS groups using the coefTest function in MATLAB.

Finally, to characterize the relationship between cortical epileptic spikes (CES) and thalamic epileptic spikes (TES), we divided the time series into 16 ms intervals (the lowest temporal resolution for epileptic spikes) and estimated the model

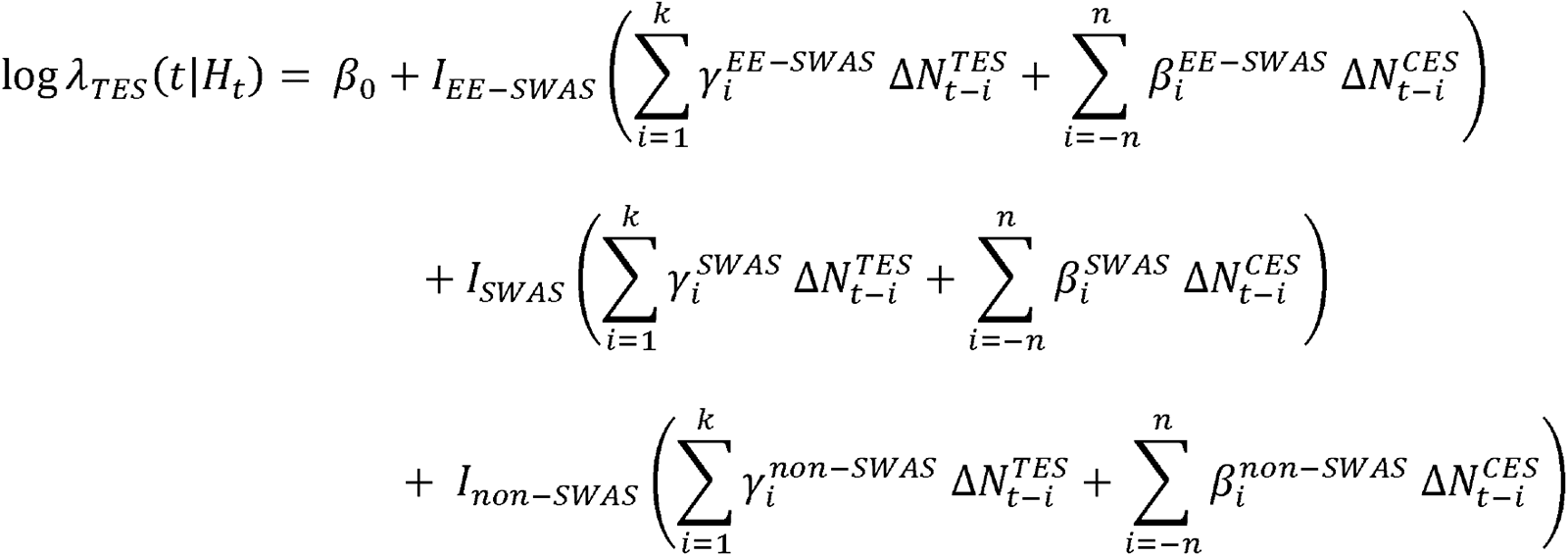

where variable *I* indicates subject group (EE-SWAS, SWAS, and non-SWAS), and we include the self-history of thalamic epileptic spikes (TES), and the history and future of cortical epileptic spikes (CES).

#### Procedures to estimate relationships between thalamic and scalp spindles

To test the hypothesis that thalamic spindles predict scalp spindles, we estimated: (1) the average evoked response at the scalp, (2) the average induced response at the thalamus, and (3) the conditional event rate of scalp spindles. We define each measure here.

To assess the average evoked response at the scalp, for each subject, we extracted from the scalp EEG signal ±2 s intervals centered on the times of the maximum amplitude of each thalamic spindle. We then estimated the mean EEG response and standard error across all subjects and thalamic spindles and tested the results against overlap with zero. The confidence limits were Bonferroni corrected for multiple comparisons.

To estimate the average induced response at the thalamus, we applied a multitaper spectral analysis to estimate the spectrogram. For each subject, we first extracted from the thalamic signal ±2 s intervals centered on the times of the maximum amplitude of each scalp spindle. We then estimated the spectrogram of each extracted thalamic signal using 0.5 s intervals (0.45 s overlap) and a single taper (2 Hz frequency resolution, function *mtspecgramc* from the Chronux toolbox (Bokil et al., 2010)). To enable testing and comparisons, we log-transformed and standardized (removed mean and divided by standard deviation) over time for each frequency of each spectrogram. We then averaged the spectrograms across all extracted signals (i.e., across all instances of scalp spindles for all subjects within each group) and tested the resulting power relative to zero using the estimated standard error.

To characterize the relationship between scalp spindles (SS) and thalamic spindles (TS), we estimated the model

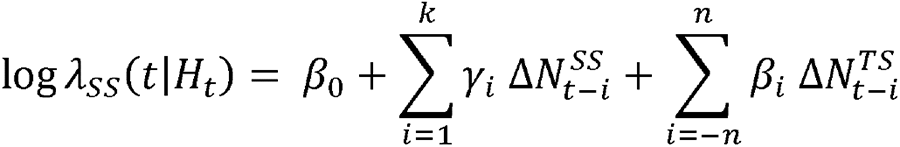

where we include the self-history of scalp spindles to account for a spindle refractory period and identify the independent contribution of thalamic spindles. We identify binary spindle occurrences using time intervals of 0.125 s, where 1 indicates the maximum amplitude of a spindle occurred (0 otherwise).

#### Procedures to estimate relationships between epileptic spikes and spindles

To test the hypothesis that epileptic spikes disrupt spindles, we examined the relationship across two timescales. At the ultradian timescale, we compared the spindle and epileptic spike rates recorded at the scalp, cortex, and thalamus. To do so, we computed for each subject the spindle and epileptic spike rates as the total number of events divided by the total duration of data analyzed (i.e., total duration of N2). We then fit a linear mixed-effects model of the spindle rates versus the log-transformed spike rates with patient as random effect. We tested the model against a nested model with constant predictor and random effect of patient using a likelihood ratio test implemented by the MATLAB function compare.

At the minutes time scale, we modeled the relationship between thalamic epileptic spikes (TES) and scalp spindles (SS) as,

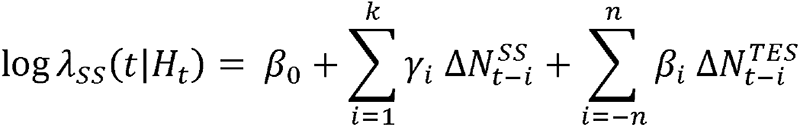

where we include the self-history of scalp spindles to account for a possible refractory period and identify the independent contribution of thalamic epileptic spikes. Here we identify binary time series for spindle events using time intervals of 1 s, where 1 indicates that the maximum amplitude of a spindle occurred in an interval. Similarly, we identify spike event time series by indicating the number of spikes (i.e., times of maximum amplitude of an epileptic spike) occurring in each 1 s time interval. We estimated the model considering self-history terms up to 20 s, and thalamic epileptic spike terms up to ±60 s (chosen based on spectrogram analysis – see Figure 4B) around each scalp spindle. Given the long timescale response observed in the data, the coefficients were tested in a group hypothesis test for the parameters between -30 s to 0 s (with 0 s representing a scalp spindle) against the parameters from 0 s to 30 s (i.e., we test whether 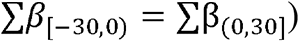.

At the seconds timescale, we compared the refractory period of scalp spindles with intervening thalamic epileptic spikes and without intervening thalamic epileptic spikes. Refractory periods were defined as the duration between the end of one spindle and the beginning of another. We fit the distributions of the refractory periods using a generalized linear model with a log-link and an inverse-Gaussian distributed response and compared the means of the two refractory period distributions using the function coefTest in MATLAB.

## Results

### Subject and data characteristics

We analyzed data from nine subjects (ages 9-55, 2 F, see clinical and data characteristics in Table 1). From each subject, voltage data were recorded from the thalamus, cortex, and scalp. Based on clinical diagnosis, and the presence spike-wave activation during N2 sleep, we separated the subjects into three categories (Supplementary Figure 1): EE-SWAS (n=3), SWAS without EE (n=3), no SWAS or EE (n=3).

**Table 1.**
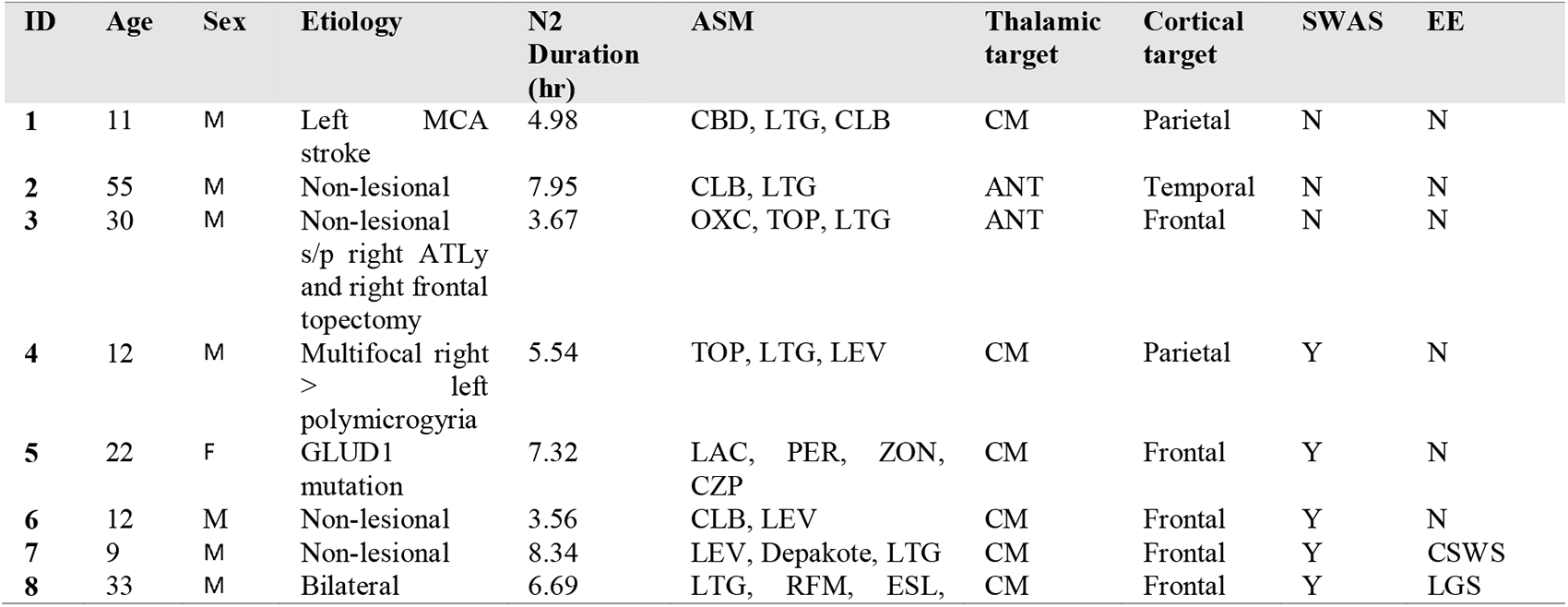
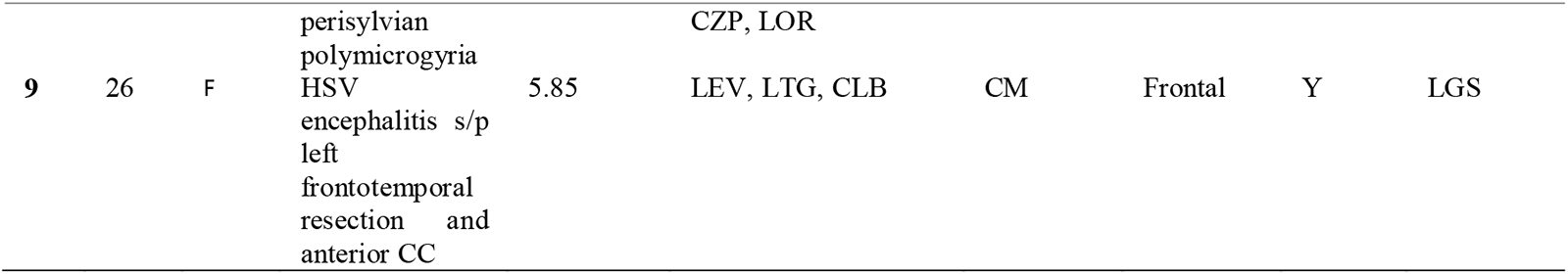
Patient Characteristics. Abbreviations: ASM-antiseizure medications; SWAS – spike and wave activation in sleep; CM – centromedian nucleus of the thalamus; ANT – anterior nucleus of the thalamus; SR-spike rate; EE-clinically diagnosed epileptic encephalopathy; CZP-clonazepam, ZON-zonisamide, PER341 perampanel, LAC-lacosamide, LTG-lamotrigine, ONFI-clobazam, LEV-levetiracetam, CBD-cannibidiol, OXC-oxcarbazepine, LOR-lorazepam, RFM-rufinamide, ESL-eslicarbazepine, ATL-anterior temporal lobectomy; LGS-lennox gastaut syndrome; CSWS-continuous spike-wave in sleep with encephalopathy, L-MCA-left middle cerebral artery, HSV-herpes simplex virus, RNS-responsive neurostimulation.

### Cortical slow oscillations facilitate spikes and spindles at different phases

To understand the influence of N2 sleep, we analyzed the relationship of slow oscillations to epileptic spikes and sleep spindles. On visual analysis, the tri-phasic slow oscillation initiates before both spikes and spindles (example in Figure 1a). Applying a statistical modeling approach to scalp slow oscillations (n=14,022), cortical epileptic spikes (n=65,107), and thalamic sleep spindles (n=14,845) (see *Methods*), we found that the first upstate of the slow oscillation began 1.4 s (defined as deviating from a baseline of 0 μV) before the down-state (i.e., at time − 1.4 s in Figure 1b). Cortical epileptic spikes were maximally coupled to the slow oscillation during the up-to-down-state transition at 0.250 s before the down-state, and sleep spindles were maximally couple to the down-to-up-state transition at 0.375 s after the down-state. Cortical spike rate was increased during the up-to-down-state transition of the slow oscillation (a positive modulation of the rate -0.75 to -0.125 s prior to the trough of a slow oscillation down-state, p<0.05 uncorrected, red curve in Figure 1b). Thalamic spindle rate was increased during the up-state (a positive modulation of the rate 0.25 s to 0.875 s after the trough of the slow oscillation down-state, p<0.05 uncorrected, green curve in Figure 1b). We conclude that cortical slow oscillations facilitate both epileptic spikes and sleep spindles, where epileptic spikes maximally occur earlier in the slow oscillation than spindles. We note that when slow oscillations are ignored, cortical spikes appear to induce spindles (Supplementary Figure 2). These results explain why epileptic spikes and spindles are both prominent in non-REM sleep, and result in the temporal correlation previously reported between epileptic spikes and spindles (Dahal et al., 2019; Gelinas et al., 2016).

**Figure 1:**
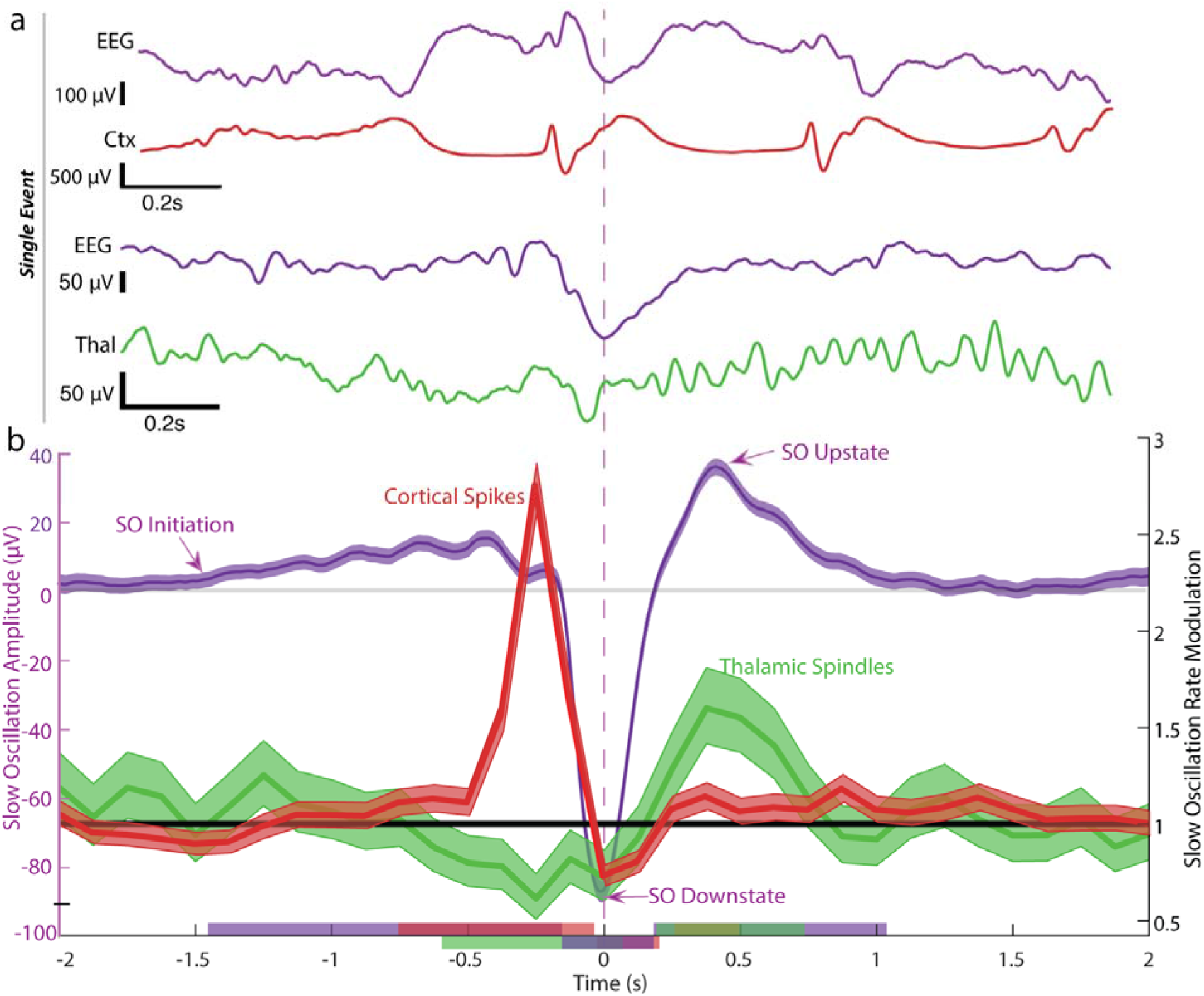
Slow waves facilitate both spikes and spindles at different phases. **(a)** Example simultaneous recordings of scalp EEG (purple trace) and intracranial data centered on the downstate of the slow oscillation (vertical dashed line). A cortical epileptiform spike (red trace) precedes the downstate and a thalamic sleep spindle (green trace) follows the downstate. **(b)** Averaged cortical slow oscillations low-pass filtered below 30 Hz (purple) across all subjects, and the modulation (95% CI) of cortical spike rate (red) and thalamic spindle rate (green) by slow waves. Slow waves initiate with a low amplitude upstate when the 95% CI exceeds 0 (left axis), approximately 1.4 s before the downstate peak. Spikes are facilitated maximally -0.5 to 0 s before the slow oscillation down state. Sleep spindles are facilitated maximally 0 to0.75 s after the slow oscillation down state (i.e., during the up-state). Significant deviations in spike rate (p<0.05) occur when the confidence limits for the modulation (right axis) do not include 1. Horizontal bars on the bottom axis indicate times of significant changes in slow oscillation amplitude (purple), spindle rate (green), and spike rate (red).

### Cortical spikes drive thalamic spikes in SWAS

Animal models suggest epileptiform spikes initiate in the cortex (Steriade & Contreras, 1998), but whether cortical epileptic spikes lead to thalamic spikes has not been evaluated (Timofeev, 2021). On visual inspection, we noted that thalamic epileptic spikes typically followed cortical epileptic spikes (example in Figure 2a). Cross-correlation histograms applied to all detected cortical epileptic spikes (n=65,107) and thalamic epileptic spikes (n=15,250) during N2 sleep revealed a similar result; thalamic spikes tended to occur after cortical spikes (Figure 2b). This relationship is most clear in subjects with EE-SWAS and SWAS, in which a higher mean number of thalamic spikes occurred within 1 s of a cortical spike compared to subjects without SWAS (F-test to compare parameters in a gamma-distributed generalized linear model, p = 4e-20, p = 6e-20, respectively; see *Methods*). To assess the relationship between cortical and thalamic spike occurrence across all subjects, we developed a generalized linear model for the time series of (binary) detected events (see *Methods*). We found that the thalamic spike rate is increased from 24 to 40 ms after a cortical spike (nested model test χ^2^(27) = 5548.1, p ≈ 0, Figure 2c). These results show that thalamic spike rates increase following a cortical spike and support the conclusion that cortical spikes drive thalamic spikes in subjects with spike-and-wave activation in sleep.

**Figure 2:**
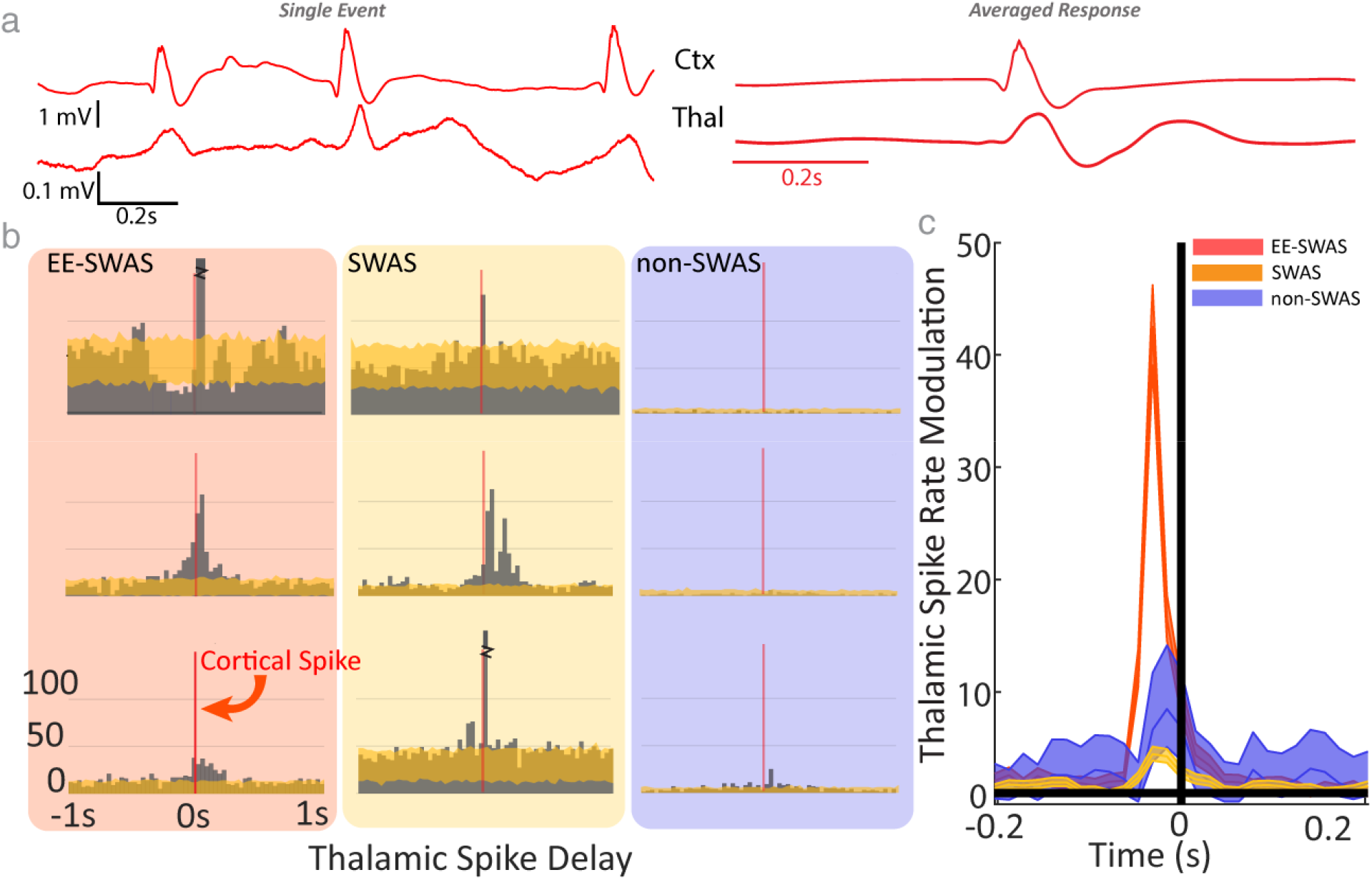
Cortical spikes drive thalamic spikes in subjects with spike and wave activation in sleep. **(a)** Example epileptic spikes (left) from one subject in the cortex (Ctx) and thalamus (Thal) and (right) the averaged response time-locked to cortical spikes. **(b)** Cross-correlation histograms of thalamic spikes relative to the time of cortical spikes for each subject. Zero indicates the moment of a cortical spike. (**c**) Model estimates of thalamic spike rate modulation due to the occurrence of a cortical spike (mean solid, confidence intervals shaded) for each patient group (see legend). Significant increases (p<0.05 when confidence intervals exclude 1) in thalamic spike rate occur due to a preceding (8 to 24 ms earlier) cortical spike.

### Spindles co-occur in thalamus and scalp EEG across all subjects

Sleep spindles are prominent rhythms generated in thalamic circuitry and transmitted to cortex, at least partially in response to cortical input (Beenhakker & Huguenard, 2009; Steriade, 2005). To confirm this relationship between thalamic and scalp spindles using direct thalamic recordings in humans, we examined the averaged scalp activity time-locked to the occurrence of thalamic spindles (see *Methods*). Thalamic spindles and scalp spindles co-occurred and were phase-locked (Figure 3a). We found higher spindle-band power in the thalamic data at the time of scalp spindles (Figure 3b) for all patient groups (standardized spectrogram power p<0.05). When modeling the scalp spindle event times using the thalamic spindle event times, we found that thalamic spindles modulated scalp spindle rate (EE-SWAS: χ^2^(20) = 138.28, p ≈ 0; SWAS: χ^2^ 2(20) = 844.52, p ≈ 0; non-SWAS: χ^2^(20) = 1230.8, p ≈ 0) across all patient groups. A thalamic spindle increased the probability of a scalp spindle by a factor of 3.3 ([2.4, 4.7] 95% CI, p=3.e-12) in EE-SWAS subjects, by a factor of 4.6 ([3.9, 5.5] 95% CI, p ≈ 0) in SWAS subjects, and by a factor of 3.1 ([2.7, 3.6] 95% CI, p ≈ 0) in non-SWAS subjects. When temporally indexed to slow oscillations (Supplementary Figure 3), the maximum modulation in the thalamic spindle rate (∼375 ms after SO downstate) preceded the maximum modulation in the scalp spindle rate (∼625 ms after SO downstate). We conclude that, across all patient groups, thalamic sleep spindles consistently drive cortical sleep spindles.

**Figure 3:**
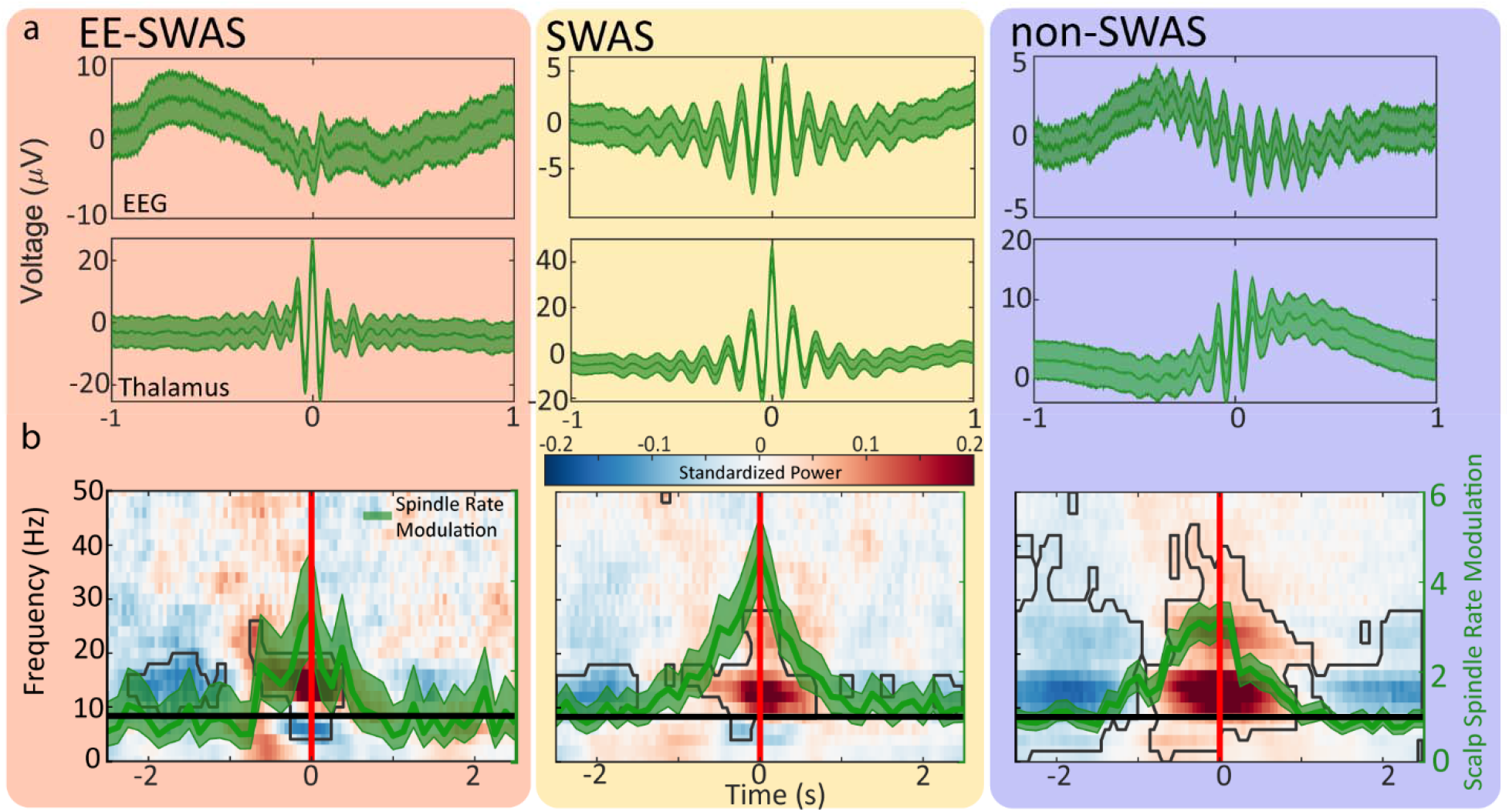
Spindles consistently co-occur in the thalamus and the scalp EEG across all subject groups. **(a)** Averaged spindle response, time-locked to maximum spindle amplitude in the thalamus, indicates consistent phase-locked spindle activity across the cortex and thalamus. **(b)** Modulation of scalp spindle rate at time 0 s (green curves, mean solid, 95% confidence intervals shaded) by thalamic spindles. Background spectrograms of thalamic data time-locked to scalp spindles. Warm (cool) colors include high (low) standardized power; see scale bar. Regions outlined in black indicate Bonferroni-corrected islands of significant changes in power.

### Spikes that propagate to the thalamus inhibit sleep spindles

While existing models suggest that epileptic spikes disrupt spindles (Beenhakker & Huguenard, 2009; Paz & Huguenard, 2015; Steriade, 2005), no direct observations of this disruption exist in humans. To address this, we evaluated the relationship between epileptic spikes and sleep spindles across scalp, cortical, and thalamic recordings. Consistent with previous observations (Kramer et al., 2021; Spencer et al., 2022), we found that the epileptic spike rate and sleep spindle rate across N2 sleep were anticorrelated (*r* = −0.59; nested model test χ^2^ (1) = 10.1, p = 0.0015; Figure 4a). We also found that, in each group, the scalp spindle refractory periods were longer if a thalamic spike occurred between spindles (EE-SWAS: 4.82 ± 0.34 s; SWAS: 3.23 ± 0.08 s; non-SWAS: 2.84 ± 0.22) compared to spindles with no intervening spike (EE-SWAS: 2.71± 0.08 s; SWAS: 1.84 ± 0.04 s; non-SWAS: 1.74 ± 0.02 s; p ≈ 0; p ≈ 0; p=1.5e-21, respectively, using an F-test for mean comparison in a generalized linear model, see Methods).

To characterize the extent of this inhibition across patient groups, we modeled the scalp spindle event rate using the history of scalp spindles and thalamic spikes as predictors (see Methods). We found additional evidence that thalamic spikes decreased spindle rate in subjects with EE-SWAS (χ^2^(121) = 216.07, p = 2e-7), but not in the other groups (SWAS: χ^2^(121) =135.46, p = 0.17; non-SWAS: χ^2^(121) = 107.91, p=0.8). For the subjects with EE-SWAS, sleep spindles were down-regulated for 30 s after a thalamic spike (*F_EE-SWAS_* (1,75010) = 43.98, p =3.4e-11; Figure 4d). Spindle rates were maximally down-regulated by a preceding (within ∼5 s) spike by a factor of 2.5 ([1.5, 4.3] 95% CI, p = 0.0011). We conclude that, across subjects, thalamic spikes increase sleep spindle refractory periods, and this mechanism decreases sleep spindle rate in subjects with EE-SWAS.

**Figure 4:**
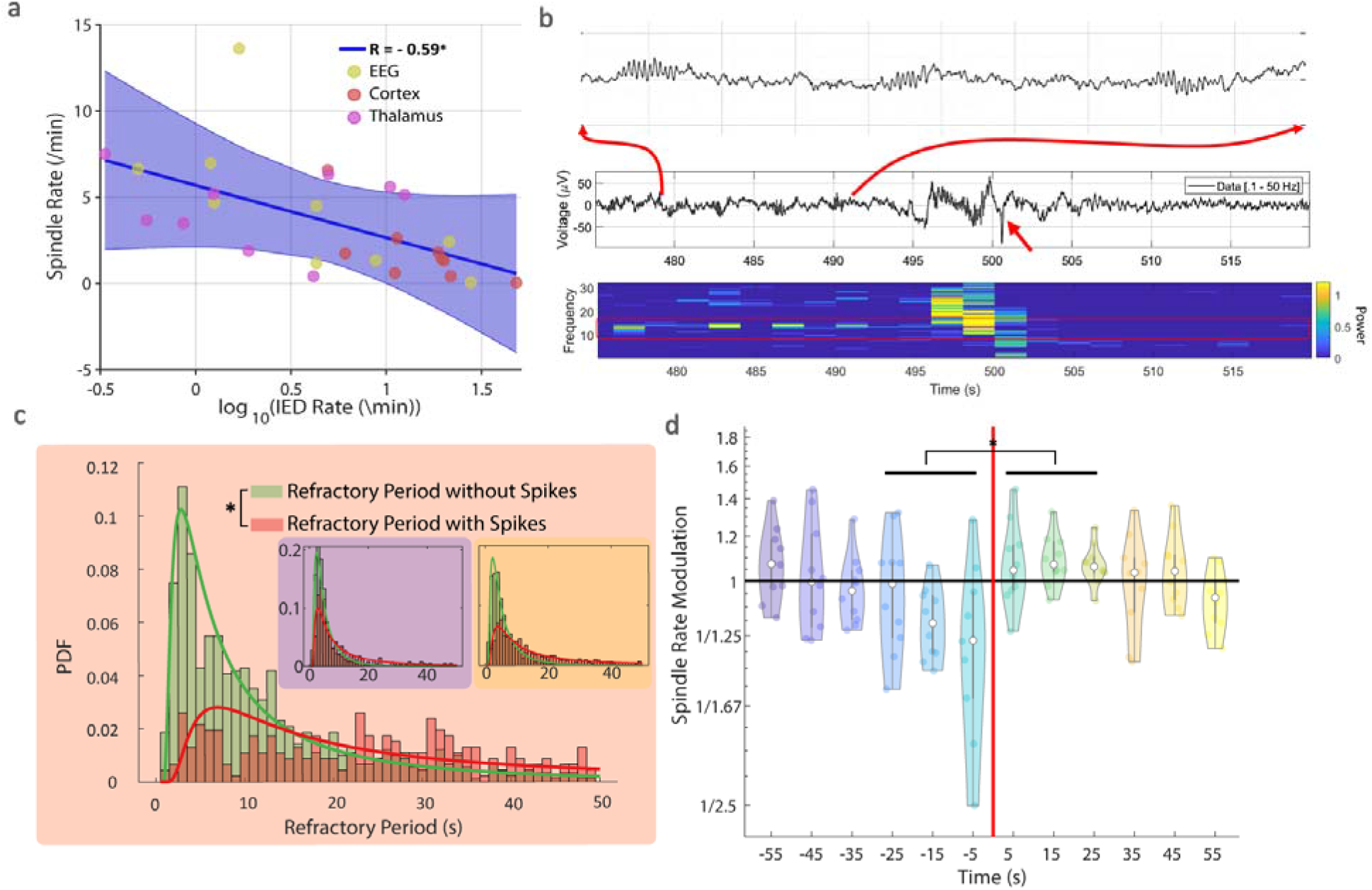
Spikes inhibit spindles. **(a)** Spike rate and spindle rate across subjects and brain regions (see legend) during N2 sleep are anti-correlated across subjects. **(b)** Example spike disruption of spindles. Before a spike (red arrow), spindles regularly occur; see upper trace for example spindles and lower image for spindle band peaks in the spectrogram. After a spike, spindles are not apparent in the trace (middle row) or spectrogram (bottom row). **(c)** Histograms of EEG spindle refractory periods with (red) and without (green) intervening thalamic spikes for subjects with epileptic encephalopathy. Insets show distributions for subjects without SWAS (blue) and with SWAS (orange). **(d)** The rate of spindles in the EEG is reduced by thalamic spikes in subjects with EE-SWAS. Estimates of spindle rate modulation parameters at each millisecond (colored dots) and grouped over 10 s intervals (violin plots with median and quartiles indicated). EEG spindle rates are down-regulated when a preceding spike occurs between -30 s to 0. * p<0.05

### Conceptual model of epileptic encephalopathy

Our analyses above reveal a unifying model of epileptic encephalopathy (Figure 5). Slow oscillations, generated in the cortex, create conditions in the brain that facilitate corticothalamic spikes and thalamocortical spindles. The up-state to down-state transition of the slow oscillation promotes pathological epileptic cortical spikes. Epileptic spikes facilitated in cortex through this process can propagate to the thalamus, a process more likely in subjects with spike and wave activation during sleep. Simultaneously, the down-state of a slow oscillation facilitates sleep spindles, which peak at the subsequent up-state of the slow oscillation. These thalamic spindles propagate to the cortex and are observable in the scalp EEG. Epileptic spikes present in the thalamus induce a longer refractory period for spindles and can result in a decrease in sleep spindle production in subjects with EE-SWAS. This spike-disrupted electrophysiological circuit demonstrates a mechanistic circuit for cognitive dysfunction in epilepsy.

**Figure 5:**
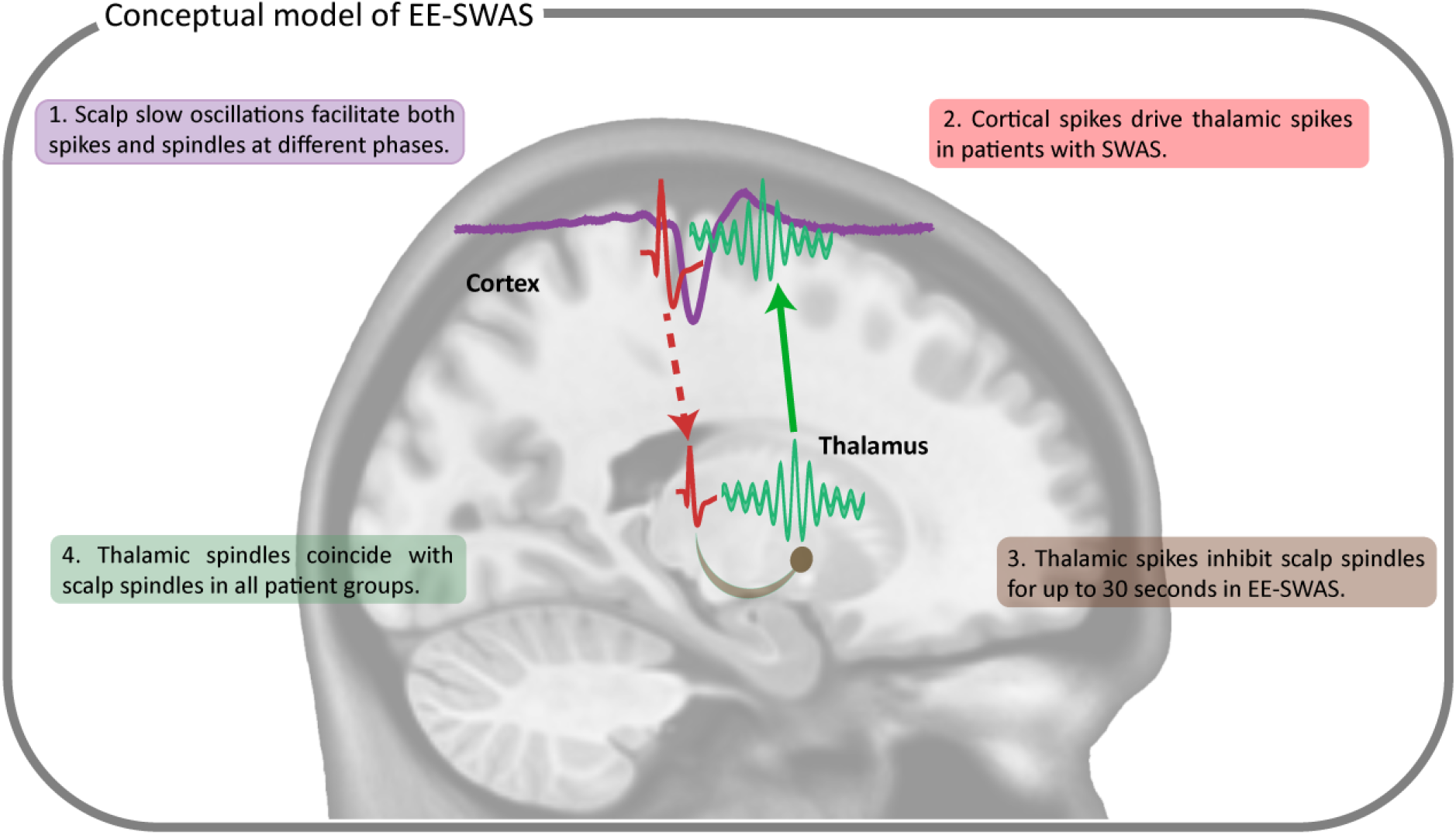
Conceptual model connecting slow oscillations, spikes, and spindles in non-REM sleep. Slow waves generated in the cortex and visible at the scalp facilitate, at different phases, both spike and spindle activity. Spikes generated in the cortex propagate to the thalamus and contribute to blocking spindles by increasing their refractory period and decreasing spindle production in subjects with EE-SWAS. Spindles generated in the thalamus propagate to the cortex in all patient groups.

## Discussion

Cognitive symptoms are a common comorbidity in epilepsy, but there are currently no mechanisms to explain cognitive dysfunction or treatments to improve them. Although an underlying etiology can often result in both epilepsy and cognitive dysfunction, and antiseizure medications themselves can contribute to cognitive dysfunction, epileptic encephalopathies are disorders in which the abnormal epileptic activity is thought to contribute directly to cognitive dysfunction beyond these influences. Here, using direct thalamic and cortical human brain recordings in subjects with epileptic activity, we reveal the thalamocortical circuitry and mechanism through which epileptic activity can contribute to cognitive dysfunction.

Slow oscillations are a ubiquitous phenomenon in non-REM sleep and have been implicated in the initiation of sleep spindles (Mak-McCully et al., 2017) and the facilitation of epileptic spikes (Frauscher et al., 2015). Inhibitory neurons may be coordinated by the slow oscillation up-state to help generate the down-state, and this increase in synchrony may pathologically facilitate an epileptic spike (see Frauscher et al., 2015 for a discussion). Alternatively, Gelinas and colleagues (Gelinas et al., 2016) suggest spikes could act as a stimulus, similar to intracortical electrical stimulation (Vyazovskiy et al., 2009), that generates a slow oscillation. In our analysis, the temporal precedence of the slow oscillation up-state contradicts the idea that the epileptic spike occurs first. We note that Gelinas and colleagues (Gelinas et al., 2016) observed alignment of delta phase in the prefrontal cortex to hippocampal epileptic spikes. In contrast, we analyzed slow oscillations at the scalp relative to cortical epileptic spikes in the epileptogenic zone, as did (Frauscher et al., 2015). This suggests the phenomenon observed by Gelinas et al. may be specific to the pre-frontal cortex given its monosynaptic connectivity to the hippocampus, while within the cortex, slow oscillations facilitate epileptic spikes in patients with epilepsy.

Across timescales and recording locations, we observed that spikes and spindles were anti-correlated. While subjects with spike activation during N2 sleep had higher overall epileptic spike rates, subjects with epileptic encephalopathies had higher likelihood of the sleep-activated cortical spikes propagating to the thalamus. Further, only subjects with epileptic encephalopathy had inhibition of sleep spindles by thalamic spikes. We conclude from these results that three separate mechanisms are required for epileptic encephalopathy – slow oscillation facilitation of epileptic spikes, corticothalamic spike propagation, and inhibition of thalamic sleep spindles. The uncovering of the thalamic neurophysiology here helps to reconcile why some subjects can have near-continuous spike and wave activity during sleep with a range of cognitive symptoms, spanning from subtle deceleration to profound cognitive regression (Wickens et al., 2017).

The role of sleep spindles in cognitive function, especially memory consolidation, in healthy individuals is well-established (Fernandez & Lüthi, 2020; Fogel & Smith, 2011; M. A. Hahn et al., 2020). Inhibition of spindles may reduce opportunities for high-fidelity memory consolidation (Dickey et al., 2021) through reactivation of neuronal patterns during hippocampal ripples (Diba & Buzsáki, 2007; Latchoumane et al., 2017). In support of this claim, spindle deficits have been linked to cognitive dysfunction in epilepsy (Kramer et al, 2021; Spencer et al 2022,Holmes & Lenck-Santini, 2006) and in response to treatment (McLaren et al., 2022). Thus, in subjects with epileptic encephalopathy, cognitive deficits may be partly explained by the reduction of sleep spindles. As sleep spindles are easily accessible in the scalp EEG, measurements of these rhythms provide a direct mechanistic biomarker of the cognitive impact of epileptic spikes. Such a biomarker could support detection of at-risk subjects, measurements of treatment response, and screening of cognitive therapeutics.

Past work suggests epileptic spikes are pathological excitatory pulses (Traub & Wong, 1982) and sleep spindle probability is regulated by activation of the thalamic reticular nucleus and the hyperpolarization of the thalamocortical neurons (Fernandez & Lüthi, 2020). Both the thalamic reticular nucleus neurons and thalamocortical neurons have a refractory period after a spindle because of after-hyperpolarization (Bal & McCormick, 1996) or after-depolarization (Kim & Mccormick, 1998), respectively. It is possible that the excess excitation induced from epileptic spikes reduces the sleep spindle probability by inducing increased depolarization of thalamocortical cells, findings consistent with a thalamocortical computational model (Li et al., 2021). Alternatively, spikes may act as de-facto spindles by occurring through the same circuits (Beenhakker & Huguenard, 2009; Steriade, 1990) and thus, enable a phase-reset to the spindle refractory period (by simultaneously inducing after-depolarization of thalamocortical cells and after-hyperpolarization of the thalamic reticular neurons).

Like most analyses of intracranial human thalamic data, our analyses were limited to a small group of subjects. Because we analyzed subjects with a wide range of ages and epilepsies, we cannot provide a comprehensive assessment of how these results vary with patient features. However, despite this diversity of patient features, we found consistent patterns across the large numbers of slow oscillations, spindles, and epileptic spikes analyzed. Future work may extend the analyses conducted here to a larger population of subjects.

Thalamic recordings are sparse and chosen based on clinical demands. As such, we combined recordings from anterior and centromedian nuclei. Recent work (Schreiner et al., 2022) has suggested that there may be differences in which area receives slow oscillations from the cortex (centromedian) and which area generates slow oscillations (anterior nucleus). In Schreiner et al. (2022) only 30% of neocortical slow oscillations were connected to anterior thalamic slow oscillations, suggesting most neocortical slow oscillations may still be cortically generated.

## Conclusion

We analyzed simultaneous invasive thalamic and cortical human recordings and non-invasive scalp EEG to understand the relationships between sleep slow oscillations, sleep spindles, and epileptic spikes. These results support a novel mechanism of cognitive dysfunction in epileptic encephalopathy, wherein epileptic spikes travel from the cortex to the thalamus, inhibit sleep spindles, and thereby disrupt the neurophysiological rhythms underlying memory consolidation.

## Notes

### Competing Interest Statement

The authors have declared no competing interest.

## References

Bal, T., & McCormick, D. A. (1996). What stops synchronized thalamocortical oscillations? Neuron, 17(2), 297–308. https://doi.org/10.1016/s0896-6273(00)80161-0

Beenhakker, M. P., & Huguenard, J. R. (2009). Neurons that Fire Together Also Conspire Together: Is Normal Sleep Circuitry Hijacked to Generate Epilepsy? Neuron, 62(5), 612– 632. https://doi.org/10.1016/j.neuron.2009.05.015

Binnie, C. D. (2003). Cognitive impairment during epileptiform discharges: Is it ever justifiable to treat the EEG? The Lancet Neurology, 2(12), 725–730. https://doi.org/10.1016/S1474-4422(03)00584-2

Bjørnæs, H., Bakke, K. A., Larsson, P. G., Heminghyt, E., Rytter, E., Brager-Larsen, L. M., & Eriksson, A.-S. (2013). Subclinical epileptiform activity in children with electrical status epilepticus during sleep: Effects on cognition and behavior before and after treatment with levetiracetam. Epilepsy & Behavior, 27(1), 40–48. https://doi.org/10.1016/j.yebeh.2012.12.007

Bokil, H., Andrews, P., Kulkarni, J. E., Mehta, S., & Mitra, P. P. (2010). Chronux: A platform for analyzing neural signals. Journal of Neuroscience Methods, 192(1), 146–151. https://doi.org/10.1016/j.jneumeth.2010.06.020

Dahal, P., Ghani, N., Flinker, A., Dugan, P., Friedman, D., Doyle, W., Devinsky, O., Khodagholy, D., & Gelinas, J. N. (2019). Interictal epileptiform discharges shape large-scale intercortical communication. Brain, 142(11), 3502–3513. https://doi.org/10.1093/brain/awz269

Denis, D., Mylonas, D., Poskanzer, C., Bursal, V., Payne, J. D., & Stickgold, R. (2021). Sleep Spindles Preferentially Consolidate Weakly Encoded Memories. Journal of Neuroscience, 41(18), 4088–4099. https://doi.org/10.1523/JNEUROSCI.0818-20.2021

Diba, K., & Buzsáki, G. (2007). Forward and reverse hippocampal place-cell sequences during ripples. Nature Neuroscience, 10(10), 1241–1242. https://doi.org/10.1038/nn1961

Dickey, C. W., Sargsyan, A., Madsen, J. R., Eskandar, E. N., Cash, S. S., & Halgren, E. (2021). Travelling spindles create necessary conditions for spike-timing-dependent plasticity in humans. Nature Communications, 12(1), Article 1. https://doi.org/10.1038/s41467-021-21298-x

Eden, U. T., & Kramer, M. A. (2010). Drawing inferences from Fano factor calculations. Journal of Neuroscience Methods, 190(1), 149–152. https://doi.org/10.1016/j.jneumeth.2010.04.012

Fernandez, L. M. J., & Lüthi, A. (2020). Sleep Spindles: Mechanisms and Functions. Physiological Reviews, 100(2), 805–868. https://doi.org/10.1152/physrev.00042.2018

Fogel, S. M., & Smith, C. T. (2011). The function of the sleep spindle: A physiological index of intelligence and a mechanism for sleep-dependent memory consolidation. Neuroscience and Biobehavioral Reviews, 35(5), 1154–1165. https://doi.org/10.1016/j.neubiorev.2010.12.003

Frauscher, B., von Ellenrieder, N., Ferrari-Marinho, T., Avoli, M., Dubeau, F., & Gotman, J. (2015). Facilitation of epileptic activity during sleep is mediated by high amplitude slow waves. Brain: A Journal of Neurology, 138(Pt 6), 1629–1641. https://doi.org/10.1093/brain/awv073

Gadot, R., Korst, G., Shofty, B., Gavvala, J. R., & Sheth, S. A. (2022). Thalamic stereoelectroencephalography in epilepsy surgery: A scoping literature review. Journal of Neurosurgery, 1–16. https://doi.org/10.3171/2022.1.JNS212613

Gais, S., Mölle, M., Helms, K., & Born, J. (2002). Learning-Dependent Increases in Sleep Spindle Density. Journal of Neuroscience, 22(15), 6830–6834. https://doi.org/10.1523/JNEUROSCI.22-15-06830.2002

Gelinas, J. N., Khodagholy, D., Thesen, T., Devinsky, O., & Buzsáki, G. (2016). Interictal epileptiform discharges induce hippocampal–cortical coupling in temporal lobe epilepsy. Nature Medicine, 22(6), 641–648. https://doi.org/10.1038/nm.4084

Greene, W. H. (2017). Econometric Analysis. Pearson Education.

Grigg Damberger Madeleine, Gozal, D., Marcus, C. L., Quan, S. F., Rosen, C. L., Chervin, R. D., Wise, M., Picchietti, D. L., Sheldon, S. H., & Iber, C. (2007). The Visual Scoring of Sleep and Arousal in Infants and Children. Journal of Clinical Sleep Medicine, 03(02), 201–240. https://doi.org/10.5664/jcsm.26819

Hahn, M. A., Heib, D., Schabus, M., Hoedlmoser, K., & Helfrich, R. F. (2020). Slow oscillation-spindle coupling predicts enhanced memory formation from childhood to adolescence. ELife, 9, e53730. https://doi.org/10.7554/eLife.53730

Hahn, M., Joechner, A.-K., Roell, J., Schabus, M., Heib, D. P., Gruber, G., Peigneux, P., & Hoedlmoser, K. (2019). Developmental changes of sleep spindles and their impact on sleep-dependent memory consolidation and general cognitive abilities: A longitudinal approach. Developmental Science, 22(1), e12706. https://doi.org/10.1111/desc.12706

Harrison, M. T., Amarasingham, A., & Kass, E. (2013). Statistical Identification of Synchronous Spiking. CRC Press, Spike Timing: Mechanisms and Function, 42.

Henin, S., Shankar, A., Borges, H., Flinker, A., Doyle, W., Friedman, D., Devinsky, O., Buzsáki, G., & Liu, A. (2021). Spatiotemporal dynamics between interictal epileptiform discharges and ripples during associative memory processing. Brain, 144(5), 1590–1602. https://doi.org/10.1093/brain/awab044

Holmes, G. L., & Lenck-Santini, P.-P. (2006). Role of interictal epileptiform abnormalities in cognitive impairment. Epilepsy & Behavior, 8(3), 504–515. https://doi.org/10.1016/j.yebeh.2005.11.014

Iber, C., Ancoli-Israel, S., Chesson, A. L., & Quan, S. (2007). The AASM Manual for the Scoring of Sleep and Associated Events: Rules, Terminology and Technical Specifications. Westchester, IL: American Academy of Sleep Medicine.

Kim, U., & Mccormick, D. A. (1998). Functional and Ionic Properties of a Slow Afterhyperpolarization in Ferret Perigeniculate Neurons In Vitro. Journal of Neurophysiology, 80(3), 1222–1235. https://doi.org/10.1152/jn.1998.80.3.1222

Klinzing, J. G., Tashiro, L., Ruf, S., Wolff, M., Born, J., & Ngo, H.-V. V. (2021). Auditory stimulation during sleep suppresses spike activity in benign epilepsy with centrotemporal spikes. Cell Reports Medicine, 2(11), 100432. https://doi.org/10.1016/j.xcrm.2021.100432

Kramer, M. A., Stoyell, S. M., Chinappen, D., Ostrowski, L. M., Spencer, E. R., Morgan, A. K., Emerton, B. C., Jing, J., Westover, M. B., Eden, U. T., Stickgold, R., Manoach, D. S., & Chu, C. J. (2021). Focal Sleep Spindle Deficits Reveal Focal Thalamocortical Dysfunction and Predict Cognitive Deficits in Sleep Activated Developmental Epilepsy. Journal of Neuroscience, 41(8), 1816–1829. https://doi.org/10.1523/JNEUROSCI.2009-20.2020

Larsson, P. G., Bakke, K. A., Bjørnæs, H., Heminghyt, E., Rytter, E., Brager-Larsen, L., & Eriksson, A.-S. (2012). The effect of levetiracetam on focal nocturnal epileptiform activity during sleep—A placebo-controlled double-blind cross-over study. Epilepsy & Behavior: E&B, 24(1), 44–48. https://doi.org/10.1016/j.yebeh.2012.02.024

Latchoumane, C.-F. V., Ngo, H.-V. V., Born, J., & Shin, H.-S. (2017). Thalamic Spindles Promote Memory Formation during Sleep through Triple Phase-Locking of Cortical, Thalamic, and Hippocampal Rhythms. Neuron, 95(2), 424–435.e6. https://doi.org/10.1016/j.neuron.2017.06.025

Li, Q., Westover, M. B., Zhang, R., & Chu, C. J. (2021). Computational Evidence for a Competitive Thalamocortical Model of Spikes and Spindle Activity in Rolandic Epilepsy. Frontiers in Computational Neuroscience, 15, 680549. https://doi.org/10.3389/fncom.2021.680549

Mak-McCully, R. A., Rolland, M., Sargsyan, A., Gonzalez, C., Magnin, M., Chauvel, P., Rey, M., Bastuji, H., & Halgren, E. (2017). Coordination of cortical and thalamic activity during non-REM sleep in humans. Nature Communications, 8(1), Article 1. https://doi.org/10.1038/ncomms15499

Mölle, M., & Born, J. (2011). Chapter 7—Slow oscillations orchestrating fast oscillations and memory consolidation. In E. J. W. Van Someren, Y. D. Van Der Werf, P. R. Roelfsema, H. D. Mansvelder, & F. H. Lopes Da Silva (Eds.), Progress in Brain Research (Vol. 193, pp. 93–110). Elsevier. https://doi.org/10.1016/B978-0-444-53839-0.00007-7

Paz, J. T., & Huguenard, J. R. (2015). Microcircuits and their interactions in epilepsy: Is the focus out of focus? Nature Neuroscience, 18(3), 351–359. https://doi.org/10.1038/nn.3950

Reynolds, C. M., Short, M. A., & Gradisar, M. (2018). Sleep spindles and cognitive performance across adolescence: A meta-analytic review. Journal of Adolescence, 66, 55–70. https://doi.org/10.1016/j.adolescence.2018.04.003

Sákovics, A., Csukly, G., Borbély, C., Virág, M., Kelemen, A., Bódizs, R., Erőss, L., & Fabó, D. (2022). Prolongation of cortical sleep spindles during hippocampal interictal epileptiform discharges in epilepsy patients. Epilepsia, n/a(n/a). https://doi.org/10.1111/epi.17337

Schreiner, T., Kaufmann, E., Noachtar, S., Mehrkens, J.-H., & Staudigl, T. (2022). The human thalamus orchestrates neocortical oscillations during NREM sleep. Nature Communications, 13(1), Article 1. https://doi.org/10.1038/s41467-022-32840-w

Specchio, N., Wirrell, E. C., Scheffer, I. E., Nabbout, R., Riney, K., Samia, P., Guerreiro, M., Gwer, S., Zuberi, S. M., Wilmshurst, J. M., Yozawitz, E., Pressler, R., Hirsch, E., Wiebe, S., Cross, H. J., Perucca, E., Moshé, S. L., Tinuper, P., & Auvin, S. (2022). International League Against Epilepsy classification and definition of epilepsy syndromes with onset in childhood: Position paper by the ILAE Task Force on Nosology and Definitions. Epilepsia, 63(6), 1398–1442. https://doi.org/10.1111/epi.17241

Spencer, E. R., Chinappen, D., Emerton, B. C., Morgan, A. K., Hämäläinen, M. S., Manoach, D. S., Eden, U. T., Kramer, M. A., & Chu, C. J. (2022). Source EEG reveals that Rolandic epilepsy is a regional epileptic encephalopathy. NeuroImage: Clinical, 33, 102956. https://doi.org/10.1016/j.nicl.2022.102956

Staresina, B. P., Bergmann, T. O., Bonnefond, M., van der Meij, R., Jensen, O., Deuker, L., Elger, C. E., Axmacher, N., & Fell, J. (2015). Hierarchical nesting of slow oscillations, spindles and ripples in the human hippocampus during sleep. Nature Neuroscience, 18(11), 1679–1686. https://doi.org/10.1038/nn.4119

Steriade, M. (1990). Spindling, Incremental Thalamocortical Responses, and Spike-Wave Epilepsy. In M. Avoli, P Gloor, G Kostopoulos, & R. Naquet (Eds.), Generalized Epilepsy: Neurobiological Approaches (pp. 161–180). Birkhäuser. https://doi.org/10.1007/978-1-4684-6767-3_12

Steriade, M. (2005). Sleep, epilepsy and thalamic reticular inhibitory neurons. Trends in Neurosciences, 28(6), 317–324. https://doi.org/10.1016/j.tins.2005.03.007

Steriade, M. (2006). Grouping of brain rhythms in corticothalamic systems. Neuroscience, 137(4), 1087–1106. https://doi.org/10.1016/j.neuroscience.2005.10.029

Steriade, M., & Contreras, D. (1998). Spike-Wave Complexes and Fast Components of Cortically Generated Seizures. I. Role of Neocortex and Thalamus. Journal of Neurophysiology. https://doi.org/10.1152/jn.1998.80.3.1439

Steriade, M., McCormick, D. A., & Sejnowski, T. J. (1993). Thalamocortical Oscillations in the Sleeping and Aroused Brain. Science, 262(5134), 679–685. https://doi.org/10.1126/science.8235588

Stoyell, S. M., Baxter, B. S., McLaren, J., Kwon, H., Chinappen, D. M., Ostrowski, L., Zhu, L., Grieco, J. A., Kramer, M. A., Morgan, A. K., Emerton, B. C., Manoach, D. S., & Chu, C. J. (2021). Diazepam induced sleep spindle increase correlates with cognitive recovery in a child with epileptic encephalopathy. BMC Neurology, 21(1), Article 1. https://doi.org/10.1186/s12883-021-02376-5

Taylor, J., & Baker, G. A. (2010). Newly diagnosed epilepsy: Cognitive outcome at 5years. Epilepsy & Behavior, 18(4), 397–403. https://doi.org/10.1016/j.yebeh.2010.05.007

Taylor, J., Kolamunnage-Dona, R., Marson, A. G., Smith, P. E. M., Aldenkamp, A. P., Baker, G. A., & SANAD study group. (2010). Patients with epilepsy: Cognitively compromised before the start of antiepileptic drug treatment? Epilepsia, 51(1), 48–56. https://doi.org/10.1111/j.1528-1167.2009.02195.x

Timofeev, I. (2021). Just a sound: A non-pharmacological treatment approach in epilepsy. Cell Reports Medicine, 2(11), 100451. https://doi.org/10.1016/j.xcrm.2021.100451

Traub, R. D., & Wong, R. K. (1982). Cellular mechanism of neuronal synchronization in epilepsy. Science (New York, N.Y.), 216(4547), 745–747. https://doi.org/10.1126/science.7079735

Truccolo, W., Eden, U. T., Fellows, M. R., Donoghue, J. P., & Brown, E. N. (2005). A Point Process Framework for Relating Neural Spiking Activity to Spiking History, Neural Ensemble, and Extrinsic Covariate Effects. Journal of Neurophysiology, 93(2), 1074– 1089. https://doi.org/10.1152/jn.00697.2004

Velasco, M., Leon, A. E.-D., Márquez, I., Brito, F., Carrillo-Ruiz, J. D., Velasco, A. L., & Velasco, F. (2002). Temporo-spatial correlations between scalp and centromedian thalamic EEG activities of stage II slow wave sleep in patients with generalized seizures of the cryptogenic Lennox–Gastaut syndrome. Clinical Neurophysiology, 113(1), 25–32. https://doi.org/10.1016/S1388-2457(01)00707-6

Velasco, M., Velasco, F., Alcalá, H., Dávila, G., & Díaz-de-León, A. E. (1991). Epileptiform EEG Activity of the Centromedian Thalamic Nuclei in Children with Intractable Generalized Seizures of the Lennox-Gastaut Syndrome. Epilepsia, 32(3), 310–321. https://doi.org/10.1111/j.1528-1157.1991.tb04657.x

Vyazovskiy, V. V., Faraguna, U., Cirelli, C., & Tononi, G. (2009). Triggering Slow Waves During NREM Sleep in the Rat by Intracortical Electrical Stimulation: Effects of Sleep/Wake History and Background Activity. Journal of Neurophysiology, 101(4), 1921–1931. https://doi.org/10.1152/jn.91157.2008

Wickens, S., Bowden, S. C., & D’Souza, W. (2017). Cognitive functioning in children with self-limited epilepsy with centrotemporal spikes: A systematic review and meta-analysis. Epilepsia, 58(10), 1673–1685. https://doi.org/10.1111/epi.13865

