## Supplementary Information for "Human thalamic recordings reveal that epileptic spikes block sleep spindle production during non-rapid eye movement sleep"

**Supplementary Figures:**

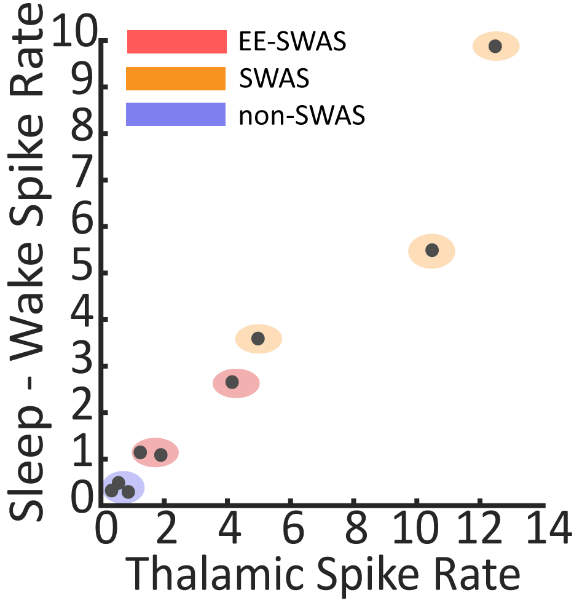

***Supp. Fig 1: Thalamic spike rates organize subjects into groups.*** *Thalamic spike rates during sleep (horizontal axis) versus the difference in cortical spike rates between wake and sleep (vertical axis). Subjects with no spike and wave activation in sleep (non-SWAS) are shown in blue, subjects clinically diagnosed with epileptic encephalopathy with spike and wave activation in sleep (EE-SWAS) in red, and the rest of the subjects are classified as having spike and wave activation in sleep (SWAS) and shown in yellow.*

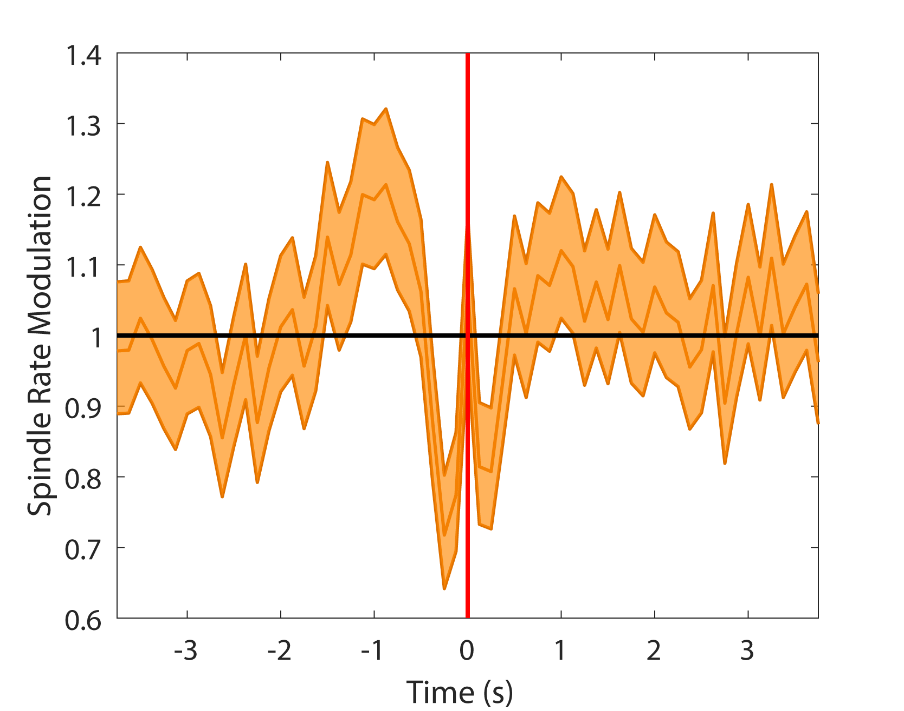

***Supp. Fig 2: Cortical spikes appear to induce thalamic spindles.*** *Model results of thalamic spindle rate with predictors cortical spikes and thalamic spindle history. The occurrence of a preceding cortical spike (near -1 s) increases the thalamic spindle rate. This temporal relationship is consistent with the relationships between spikes and spindles to slow* oscillations*. The down-regulation of spindles near 0 s indicates that spikes and spindles are less likely to co-occur.*

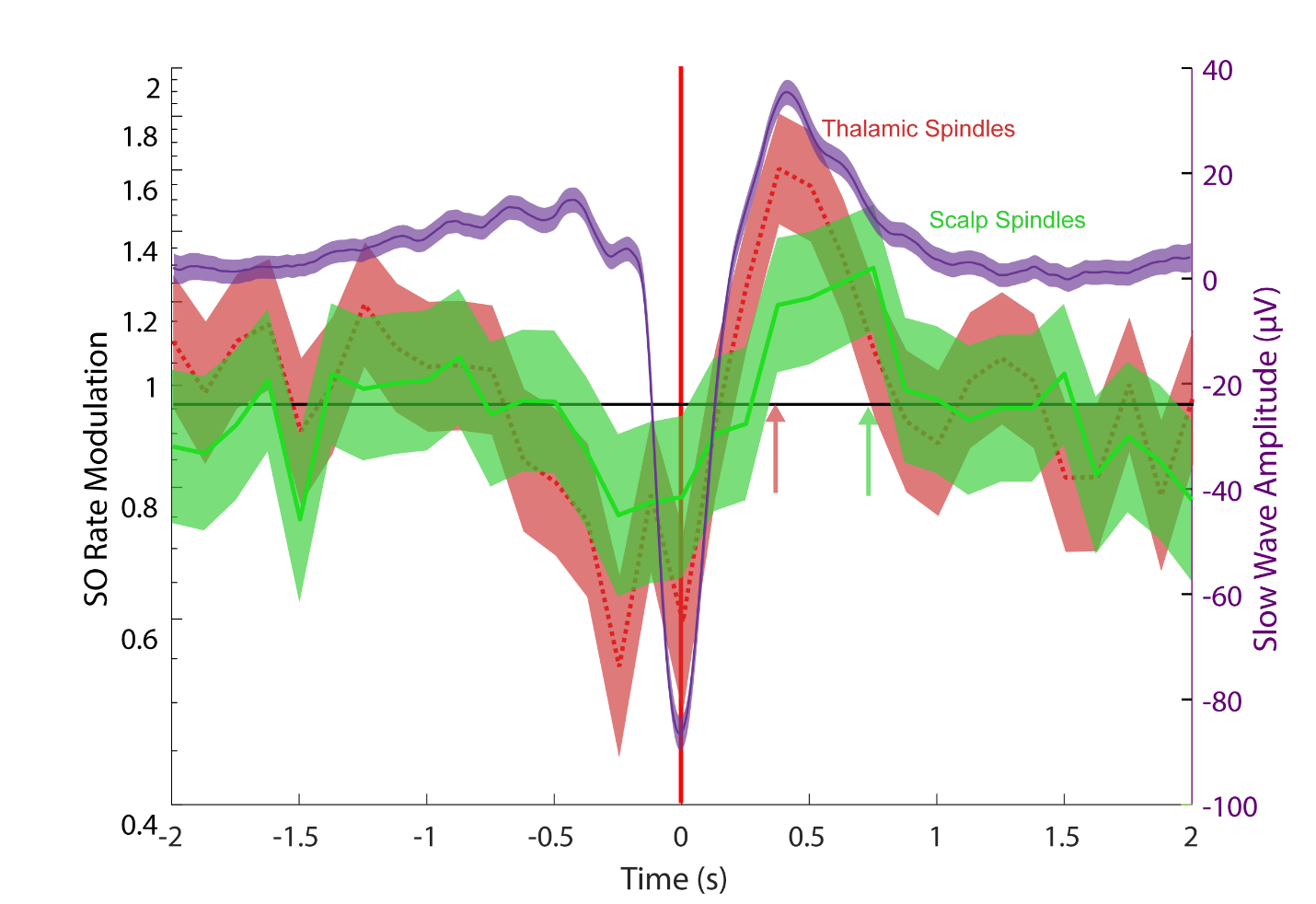

***Supp. Fig 3:*** ***Thalamic spindles precede scalp spindles when indexed to slow oscillation*** ***down-states.*** *Parameter values and confidence intervals for models of thalamic (red) and scalp (green) spindle rates with predictor slow oscillation down-state, and the average amplitude of the slow oscillation down-state (purple). Red arrow indicates time of maximum up-regulation of thalamic spindles and green arrow indicates time of maximum up-regulation of EEG spindles.*
